# Boundaries support specific long-distance interactions between enhancers and promoters in *Drosophila Bithorax* complex

**DOI:** 10.1101/423103

**Authors:** Nikolay Postika, Mario Metzler, Markus Affolter, Martin Müller, Paul Schedl, Pavel Georgiev, Olga Kyrchanova

**Author notes:** Corresponding authors: (OK), (PG).

## Abstract

*Drosophila* bithorax complex (BX-C) is one of the best model systems for studying the role of boundaries (insulators) in gene regulation. Expression of three homeotic genes, *Ubx, abd-A,* and *Abd-B*, is orchestrated by nine parasegment-specific regulatory domains. These domains are flanked by boundary elements, which function to block crosstalk between adjacent domains, ensuring that they can act autonomously. Paradoxically, seven of the BX-C regulatory domains are separated from their gene target by at least one boundary, and must “jump over” the intervening boundaries. To understand the jumping mechanism, the *Mcp* boundary was replaced with *Fab-7* and *Fab-8*. *Mcp* is located between the *iab-4* and *iab-5* domains, and defines the border between the set of regulatory domains controlling *abd-A* and *Abd-B*. When *Mcp* is replaced by *Fab-7* or *Fab-8,* they direct the *iab-4* domain (which regulates *abd-A*) to inappropriately activate *Abd-B* in abdominal segment A4. For the *Fab-8* replacement, ectopic induction was only observed when it was inserted in the same orientation as the endogenous *Fab-8* boundary. A similar orientation dependence for bypass activity was observed when *Fab-7* was replaced by *Fab-8*. Thus, boundaries perform two opposite functions in the context of BX-C – they block crosstalk between neighboring regulatory domains, but at the same time actively facilitate long distance communication between the regulatory domains and their respective target genes.

**Author Summary:** *Drosophila* bithorax complex (BX-C) is one of a few examples demonstrating *in vivo* role of boundary/insulator elements in organization of independent chromatin domains. BX-C contains three *HOX* genes, whose parasegment-specific pattern is controlled by *cis*-regulatory domains flanked by boundary/insulator elements. Since the boundaries ensure autonomy of adjacent domains, the presence of these elements poses a paradox: how do the domains bypass the intervening boundaries and contact their proper regulatory targets? According to the textbook model, BX-C regulatory domains are able to bypass boundaries because they harbor special promoter targeting sequences. However, contrary to this model, we show here that the boundaries themselves play an active role in directing regulatory domains to their appropriate *HOX* gene promoter.

## Introduction

The three homeotic (HOX) genes in the *Drosophila* Bithorax complex (BX-C), *Ultrabithorax* (*Ubx*), *abdominal-A* (*abd-A*) and *Abdominal-B* (*Abd-B*), are responsible for specifying cell identity in parasegments (PS) 5-14, which form the posterior half of the thorax and all of the abdominal segments of the adult fly [1–3]. Parasegment identity is determined by the precise expression pattern of the relevant HOX gene and this depends upon a large *cis*-regulatory region that spans 300 kb and is subdivided into nine PS domains that are aligned in the same order as the body segments in which they operate [4–6]. *Ubx* expression in PS5 and PS6 is directed by *abx/bx* and *bxd/pbx*, while *abd-A* expression in PS7, PS8, and PS9 is controlled by *iab-2, iab-3*, and *iab-4* [7–10]. *Abd-B* is regulated by four domains, *iab-5, iab-6, iab-7* and *iab-8*, which control expression in PS10, PS11, PS12 and PS13 respectively [6,11,12].

Each regulatory domain contains an initiator element, a set of tissue-specific enhancers and Polycomb Responsible Elements (PREs) and is flanked by boundary/insulator elements (Fig 1A; Maeda and Karch 2006). *BX-C* regulation is divided into two phases, initiation and maintenance [15,16]. During the initiation phase, a combination of gap and pair-rule proteins interact with initiation elements in each regulatory domain, setting the domain in the *on* or *off* state. In PS10, for example, the *iab-5* domain, which regulates *Abd-B*, is activated by its initiator element, while the more distal *Abd-B* domains, *iab-6* to *iab-8* are set in the *off* state (Fig 1B). In PS11, the *iab-6* initiator activates the domain, while the adjacent *iab-7* and *iab-8* domains are set in the *off* state. Once the gap and pair-rule gene proteins disappear during gastrulation, the *on* and *off* states of the regulatory domains are maintained by Trithorax (Trx) and Polycomb (PcG) group proteins, respectively [17,18].

**Fig 1.**
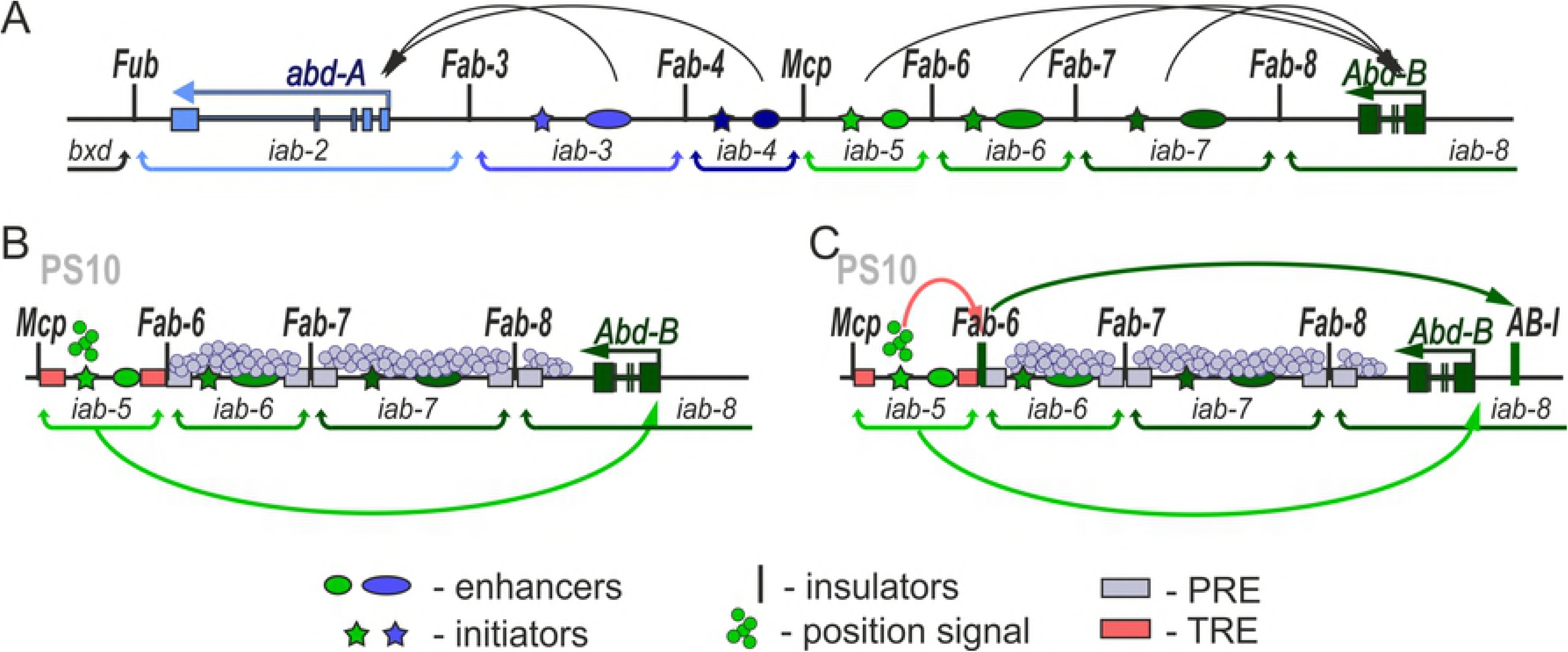
Models of an enhancer – promoter interactions in *BX-C*. (A) Regulatory region of the distal part of the BX-C. Horizontal arrows represent transcripts for *abd-A* (blue) and *Abd-B* (green). *iab* enhancers are shown as ovals color-coded with respect to the gene they control (darker shades of color indicate higher expression levels). The arrow arches are a graphical illustration of the targeting of each *cis-*regulatory domain to the *abd-A* or *Abd-Bm* promoter. Vertical lines mark boundaries (*Fub, Fab-3, Fab-4, Mcp, Fab-6, Fab-7*, and *Fab-8*) of regulatory *iab* domains which are delimited by brackets behind the map. There is also a boundary-like element *AB-I* upstream of the *Abd-B* promoter that has communicator activity in bypass assays. (B) and (C) Schematic representation of the models explaining interaction of the *iab* enhancers with the *Abd-B* promoter.

In order to select and then maintain their activity states independent of outside influence, adjacent regulatory domains are separated from each other by boundary elements or insulators [19–25]. Mutations that impair boundary function permit crosstalk between positive and negative regulatory elements in adjacent domains and this leads to the misspecification of parasegment identity. This has been observed for deletions that remove five of the BX-C boundaries (*Front-ultraabdominal* (*Fub*), *Miscadestral pigmentation* (*Mcp*), *Frontadominal-6* (*Fab-6*), *Frontadominal-7* (*Fab-7*), and *Frontadominal-8* (*Fab-8*)) [6,18,20,21,23,24,26,27].

While these findings indicate that boundaries are needed to ensure the functional autonomy of the regulatory domains, their presence also poses a paradox [14,28]. Seven of the nine BX-C regulatory domains are separated from their target HOX gene by at least one intervening boundary element. For example, the *iab-6* regulatory domain must “jump over” or “bypass” *Fab-7* and *Fab-8* in order to interact with the *Abd-B* promoter (Fig 1A). That the blocking function of boundaries could pose a significant problem has been demonstrated by experiments in which *Fab-7* is replaced by heterologous elements such as *scs, gypsy* or multimerized binding sites for the architectural proteins dCTCF, Pita or Su(Hw) [26,29–31]. In these replacements, the heterologous boundary blocked crosstalk between *iab-6* and *iab-7* just like the endogenous boundary, *Fab-7*. However, the boundaries were not permissive for bypass, preventing *iab-6* from regulating *Abd-B*.

A number of models have been proposed to account for this paradox. One is that BX-C boundaries must have unique properties that distinguish them from generic fly boundaries. Since they function to block crosstalk between enhancers and silencers in adjacent domains, an appealing idea is that they would be permissive for enhancer/silencer interactions with promoters (Fig 1B). However, several findings argue against this notion. For one, BX-C boundaries resemble those elsewhere in the genome in that they contain binding sites for architectural proteins such as Pita, dCTCF, and Su(Hw) [24,31–35]. Consistent with their utilization of these generic architectural proteins, when placed between enhancers (or silencers) and a reporter gene, BX-C boundaries block regulatory interactions just like boundaries from elsewhere in the genome [20,36–42]. Similarly, there is no indication in these transgene assays that the blocking activity of BX-C boundaries are subject to parasegmental regulation. Also arguing against the idea that BX-C boundaries have unique properties, the *Mcp* boundary, which is located between *iab-4* and *iab-5*, is unable to replace *Fab-7* [31]. Like the heterologous boundaries, it blocks crosstalk, but it is not permissive for bypass. A second model is that there are special sequences, called promoter targeting sequence (PTS), located in each regulatory domain that actively mediate bypass (Zhou and Levine 1999; Chen et al. 2005; Lin et al. 2003). While the PTS sequences that have been identified in *iab-6* and *iab-7* enable enhancers to “jump over” an intervening boundary in transgene assays, they do not have a similar function in the context of BX-C and are completely dispensable for *Abd-B* regulation [19,30].

A third model (Fig 1C) is suggested by transgene “insulator bypass” assays [46,47]. In one version of this assay, two boundaries instead of one are placed in between an enhancer and the reporter. When the two boundaries pair with each other, the enhancer is brought in close proximity to the reporter, thereby activating rather than blocking expression. Consistent with a possible role in BX-C bypass, these pairing interactions can occur over large distances and even skip over many intervening boundaries [48–51]. The transgene assays point to two important features of boundary pairing interactions that are likely to be relevant in BX-C. First, pairing interactions are specific. For this reason boundaries must be properly matched with their neighborhood in order to function appropriately. A requirement for matching is illustrated in transgene bypass experiments in which multimerized binding sites for specific architectural proteins are paired with themselves or with each other[52]. Bypass was observed when multimerized dCTCF, Zw5 or Su(Hw) binding sites were paired with themselves; however, heterologous combinations (e.g. dCTCF sites with Su(Hw) sites) did not support bypass.

The fact that both blocking and bypass activities are intrinsic properties of fly boundaries suggests that the BX-C boundaries themselves may facilitate contacts between the regulatory domains and their target genes (Fig 1C). Moreover, the non-autonomy of both blocking and bypass activity could potentially explain why heterologous *Fab-7* replacements like *gyspy* and *Mcp* behave anomalously while *Fab-8* functions appropriately. Several observations fit with this idea. There is an extensive region upstream of the *Abd-B* promoter that has been implicated in tethering the *Abd-B* regulatory domains to the promoter [53–56] and this region could play an important role in mediating bypass by boundaries associated with the distal *Abd-B* regulatory domains (*iab-5, iab-6, iab-7*). Included in this region is a promoter tethering element (PTE) that facilitates interactions between *iab* enhancers and the *Abd-B* promoter in transgene assays [57,58]. Just beyond the PTE is a boundary element, *AB-I*. In transgene assays *AB-I* mediates bypass when combined with either *Fab-7* or *Fab-8*. In contrast, a combination between *AB-I* and *Mcp* fails to support bypass [59,60]. The ability of both *Fab-7* and *Fab-8* to pair with *AB-I* is recapitulated in *Fab-7* replacement experiments. Unlike *Mcp, Fab-8* has both blocking and bypass activity when inserted into *Fab-7* [30]. Moreover, its’ bypass but not blocking activity is orientation-dependent. When inserted in the same orientation as the endogenous *Fab-8* boundary, it mediates blocking and bypass, while it does not support bypass when inserted in the opposite orientation.

In the studies reported here we have tested this model by replacing the endogenous *Mcp* boundary with heterologous boundaries. *Mcp* defines the border between the set of regulatory domains that control *abd-A* and those that control *Abd-B* expression (Fig 1A). Unlike the boundaries that are within the *Abd-B* regulatory domain (e.g. *Fab-7* or *Fab-8*), *Mcp* is not located between a regulatory domain and its target gene. Instead, it defines the boundary between regulatory domains that target *abd-A* and those that target *Abd-B*. For this reason, we expected that it does not need bypass activity. Consistent with this expectation, we find that multimerized dCTCF binding sites fully substitute for *Mcp*. A different result is obtained for the *Abd-B*-associated boundaries, *Fab-7* and *Fab-8*. Both boundaries are (for the most part) able to block crosstalk between the *abd-A* regulatory domain *iab-4*, which specifies A4 (PS9) and the *Abd-B* regulatory domain *iab-5*, which specifies A5 (PS10). Their blocking activity is orientation independent. However, in spite of blocking crosstalk, these replacements still inappropriately induce *Abd-B* expression in A4 (PS9), causing the misspecification of this segment. For the *Fab-7* replacements, this occurred in both orientations, while for the *Fab-8* replacement ectopic induction was only observed when it was inserted in the same orientation as the endogenous *Fab-8* boundary. We present evidence showing that the boundary replacements activate the *Abd-B* gene in A4 (PS9) by inappropriately targeting the *iab-4* domain to the *Abd-B* promoter. In addition to altering the specification of A4 (PS9), the *Fab-7* replacements induce novel transformations of A5 and A6. These findings indicate that when *Fab-7* is inserted into the BX-C in place of *Mcp,* it perturbs the function not only of *iab-4*, but also *iab-5* and *iab-6*.

## Results & Discussion

### Substitution of *Mcp* by an *attP* integration site in the BX-C

The *Mcp* boundary is defined by 340 bp core sequence that has enhancer blocking activity in transgene assays [36] and blocks crosstalk between *iab-6* and *iab-7* when substituted for *Fab-7* [31]. Located just distal to the boundary is a PRE that negatively regulates the activity of the *iab-5* enhancers [61]. We used CRISPR to delete a 789 bp DNA segment including the *Mcp* boundary and the PRE and replace it with an *eGFP* reporter flanked by two *attP* sites (*Mcp*^*attP*^) (S1 Fig). The presence of two *attP* sites in opposite orientation allows integration of different DNA fragments by recombination mediated cassette exchange (RMCE; Bateman et al) using the *phiC31* integration system [62].

### Multimerized dCTCF sites substitute for *Mcp*

The *Mcp* boundary marks the division between the set of regulatory domains that control the *abd-A* and *Abd-B* genes (Fig 1A). The *iab-4* domain is just proximal to *Mcp*, and it directs *abd-A* expression in PS9. The *iab-5* domain is on the distal side and it regulates *Abd-B* in PS10. A boundary in this position would be needed to block crosstalk between *iab-4* and *iab-5*; however, neither of these domains would require the intervening boundary to have bypass activity. On the proximal side, *iab-4* must bypass the putative *Fab-3* and *Fab-4* in order to activate the *abd-A* promoter, while on the distal side, *iab-5* must bypass *Fab-6, Fab-7* and *Fab-8* in order to activate *Abd-B*. If this expectation is correct, a generic boundary that has blocking activity but is unable to direct *iab-4* to the *abd-A* promoter or *iab-5* to the *Abd-B* promoter should be able to substitute for *Mcp*. To test this prediction (Fig 2), we introduced either the *iab-5* PRE itself (*Mcp*^*PRE*^) or the PRE in combination with four dCTCF sites (*Mcp*^*CTCF*^). In *Fab-7* replacement experiments four dCTCF sites in combination with the *iab-7* PRE block crosstalk between the *iab-6* and *iab-7* domains, but do not allow the *iab-6* domain to regulate *Abd-B* expression in PS11 [30].

**Fig 2.**
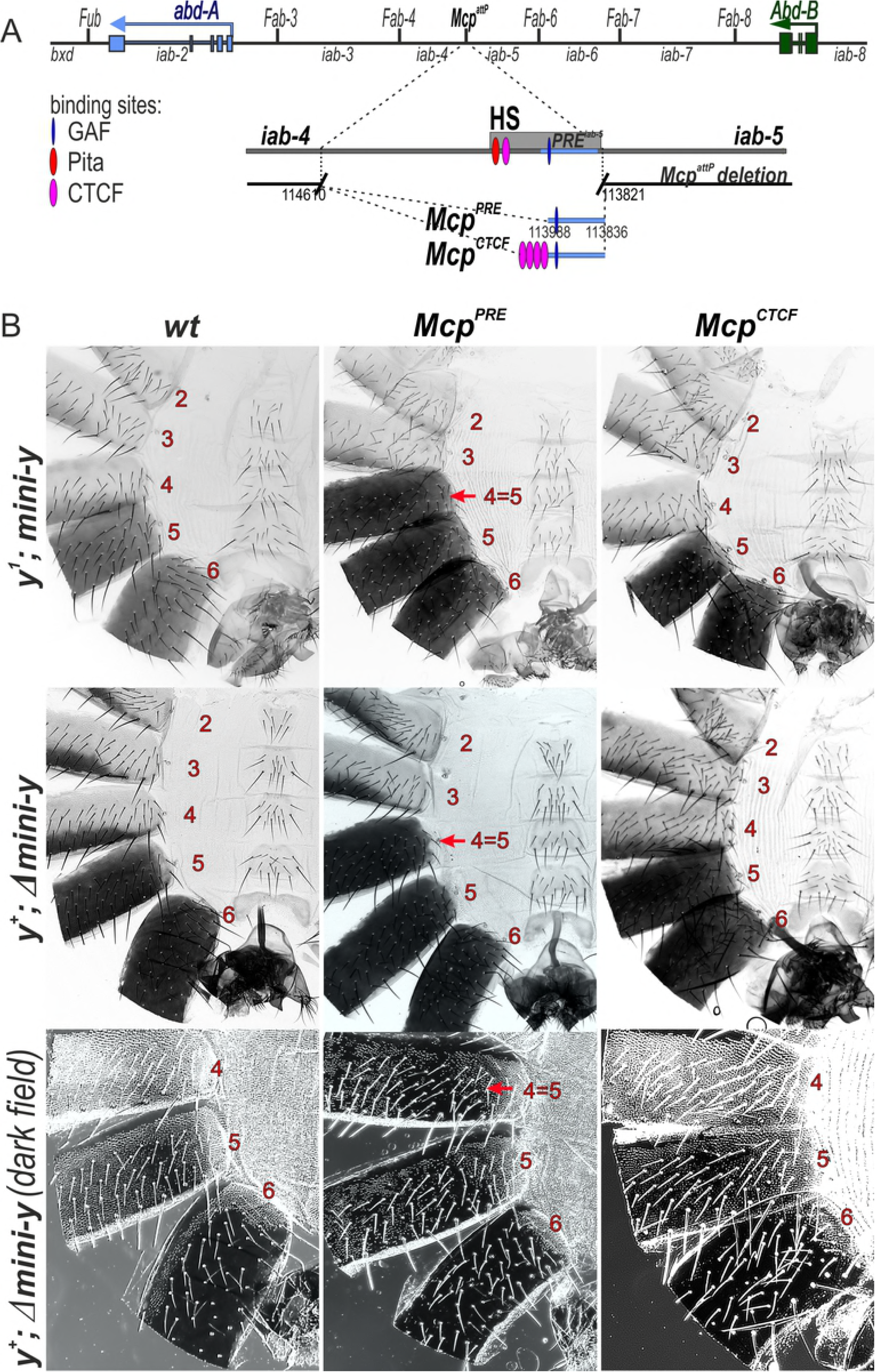
The CTCF sites block crosstalk between the *iab-4* and *iab-5* domains. (A) Molecular maps of the *Mcp* boundary. The coordinates of the *Mcp*^*attP*^ deletion and *Mcp*^*PRE*^ *Mcp*^*CTCF*^ replacement fragments according to complete sequence of BX-C in SEQ89E numbering [4] are shown below. DNAse hypersensitive site is shown as a light gray box above the coordinate bar. Binding sites for GAF, Pita and dCTCF are indicated by blue, red and green ovals, respectively. PRE element from *iab-5* is marked as a blue stripe. Replacement fragments are shown below. (B) The cuticle preparations of *wt, Mcp*^*PRE*^ and *Mcp*^*CTCF*^ males. The morphology of the 2th to 6th abdominal segments is shown. Abnormalities in segment phenotype are shown by the red arrows. The localization of trichomes on the 4th to 6th abdominal tergites are shown in dark field.

The *Abd-B* protein is master regulator of pigmentation in the male abdominal A5 and A6 segments due to the regulation of genes involved in melanin synthesis [63–65]. Flies carrying the null *y*^*1*^ allele lack black melanin but still have brown melanin that is also regulated by the Abd-B protein [65,66]. In order to be able to recover recombinants and also to monitor the blocking activity of the replacement sequence and the *on*/*off* state of the *iab-5* domain, we included a minimal *yellow* (*mini-y*) reporter in our *Mcp* replacement construct. The *mini-y* reporter consists of the cDNA and the 340 bp *yellow* promoter and lacks the wing, body and bristle enhancers of the endogenous *yellow* gene. As a result, activity of the *mini-y* reporter depends upon proximity to nearby enhancers. Expression of the *mini-y* reporter was examined in the *y*^*1*^ background.

Based on previous studies [5,22,67], we assume that the activity of this reporter will be determined by the activity state of the *iab-5* domain. When *iab-5* is *off* in PS9 and more anterior parasegments, the *mini*-*y* reporter will also be off. When *iab-5* is *on* in PS10 and more posterior parasegments, the *mini*-*y* reporter will be expressed. This parasegment-specific regulation of the reporter activity will be reflected in the segmental pattern of black melanin pigmentation in the adult cuticle. In replacements in which blocking activity is compromised, *mini*-*y* will be expressed in PS9 in adults the A4 tergite will be black, just like the A5 and A6 tergites.

When we replaced the *Mcp* deletion by the *iab-5* PRE alone (*Mcp*^*PRE*^) the *mini*-*y* reporter was active not only in A5 (PS10) and more posterior segments, but also in A4 (PS9). As shown in Fig 2, the pigmentation in A4 is black like that in A5 indicating that the reporter is expressed in both segments (Fig 2). This finding shows that, similar to classical *Mcp* deletions, the *Mcp*^*PRE*^ replacement does not have blocking activity. In these *Mcp* deletions *iab-5* is ectopically activated in PS9 by the *iab-4* initiator and as a consequence there is a gain-of-function transformation in parasegment identity from PS9 to PS10. We used two approaches to test whether this was true for the *Mcp*^*PRE*^ replacement. In the first, we excised the *mini-y* reporter and introduced an *y*^*+*^ X chromosome. Since *Abd-B* directly regulates endogenous *yellow* expression in abdomen [64,66], a transformation of PS9 into PS10 should be accompanied by a PS10-like pattern of pigmentation. Fig 2 shows that this is indeed the case. We also examined the pattern of Abd-B protein expression in the embryonic CNS. In wild type embryos Abd-B is not expressed is PS9, while it is expressed at low levels in PS10. As shown in Fig 3A, Abd-B protein is detected in both PS9 and PS10 at similarly high levels in the *Mcp*^*PRE*^ replacement.

**Fig 3.**
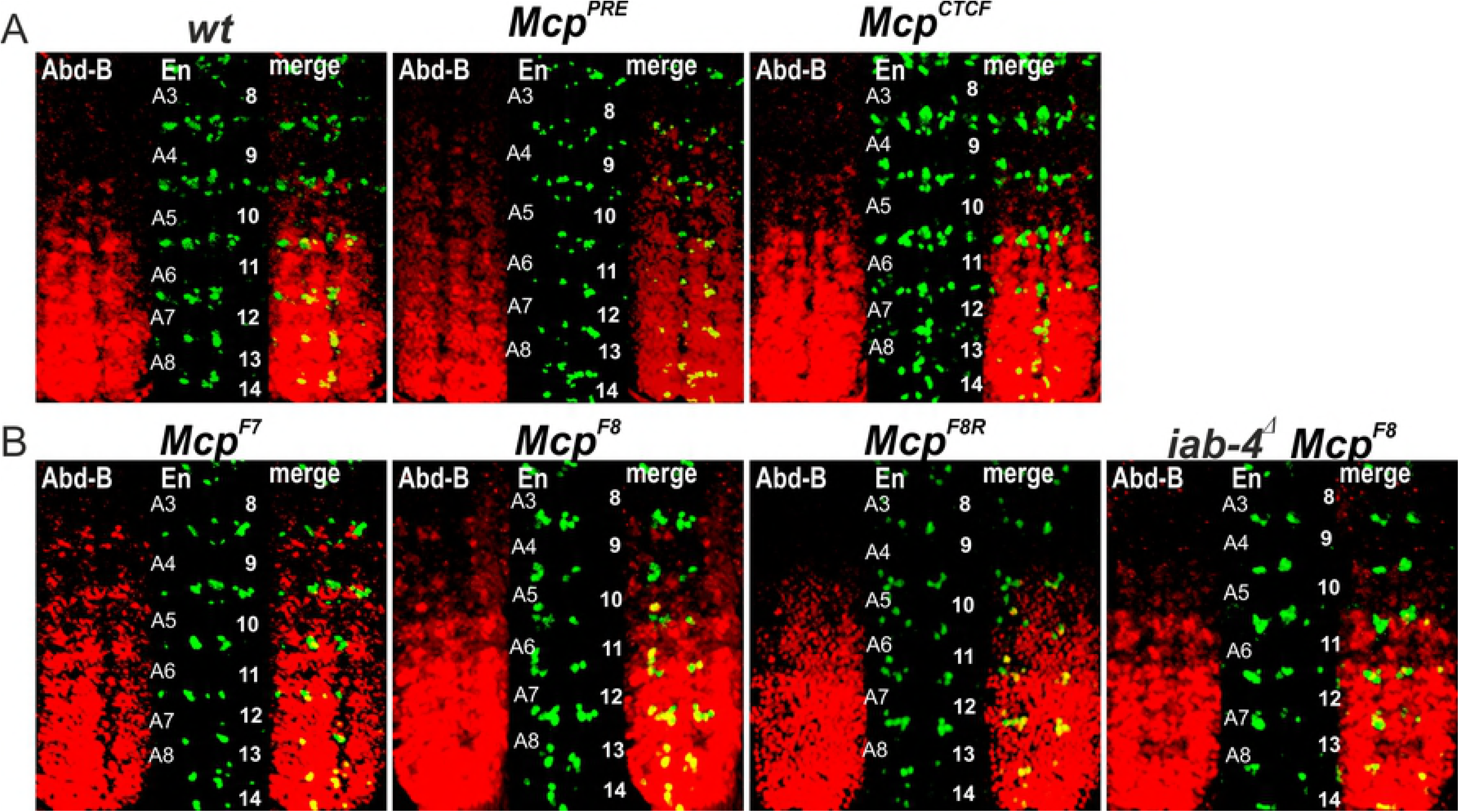
Expression of *Abd-B* in *Mcp* replacement embryos. (A) *Abd-B* expression in *wt, Mcp*^*PRE*^ and *Mcp*^*CTCF*^ embryos. (B) *Abd-B* expression in *Mcp*^*F7*^, *Mcp*^*F8*^, *Mcp*^*F8R*^ and *iab-4*^*Δ*^ *Mcp*^*F8*^ embryos. Each panel shows an image of the embryonic CNS of stage 14 embryos stained with antibodies to ABD-B (red) and Engrailed (En, green). En is used to mark parasegments, which are numbered from 9 to 14 on the right side of the panels; approximate positions of segments are shown on the left side of the wild type (wt) panel and marked A4 to A8. The wild type expression pattern of *Abd-B* in the embryonic CNS is characterized by a stepwise gradient of increasing protein level from PS10 to PS14. The *Mcp*^*F8*^ or *Mcp*^*F7*^ embryos have similar low *Abd-B* expression in PS9 and PS10. The *Abd-B* expression in PS9 is absent in *iab-4*^*Δ*^ *Mcp*^*F8*^ and *Mcp*^*F8R*^ embryos.

As predicted, a quite different result is obtained when we combined the *iab-5* PRE with multimerized dCTCF sites. Expression of the *mini-y* reporter in the *Mcp*^*CTCF*^ replacement was restricted to A5 (PS10) and A6 (PS11) as would be expected if the multimerized dCTCF sites block crosstalk between the *iab-4* and *iab-5* domains (Fig 2). The same pigmentation pattern is observed for the endogenous *yellow* in the Δ*mini-y* derivative of *Mcp*^*CTCF*^, indicating that *Abd-B* is not turned on ectopically in PS9. This conclusion is confirmed by antibody staining experiments (Fig 3A). Thus, unlike replacements of *Fab-7*, a generic boundary can fully substitute for *Mcp*.

### Substitution of *Fab-7* for *Mcp* disrupts *Abd-B* regulation in parasegments PS9, PS10 and PS11

We next tested whether the *Fab-7* boundary can substitute for *Mcp*. The *Fab-7* region consists of a minor (HS*) and three major (HS1, HS2 and HS3) nuclease hypersensitive sequences [18,22,23,41,42]. Unlike *Mcp* or other known or suspected boundaries in BX-C, dCTCF does not bind to *Fab-7* [33,68]. Instead, *Fab-7* boundary function depends upon two BEN domain protein complexes, Elba and Insensitive, the C_2_H_2_ zinc finger protein Pita, and a large multiprotein complex, called the LBC (Fedotova et al. 2017; Kyrchanova et al. 2017; Wolle et al. 2015; Cleard et al. 2017; Kyrchanova et al. 2018). In addition to a boundary function, the HS3 sequence can also function as a PRE (*iab-7* PRE; Mishra et al. 2001; Kyrchanova et al. 2018). In previous studies, we found that a combination of HS1+HS2+HS3 can functionally substitute for the complete *Fab-7* boundary *in vivo* and we used this sequence (named for simplicity *F7*) for the *Mcp* replacements (Fig 4). Although *Fab-7* has only limited orientation dependence in its endogenous context (Kyrchanova et al. 2016; Kyrchanova et al. 2018), we inserted the HS1+HS2+HS3 sequence in both the forward (same as endogenous *Fab-7*) and reverse orientations in the *Mcp* replacement platform. The phenotypic effects of the *Fab-7* replacement inserted in the forward orientation, *Mcp*^*F7*^, are considered first.

**Fig 4.**
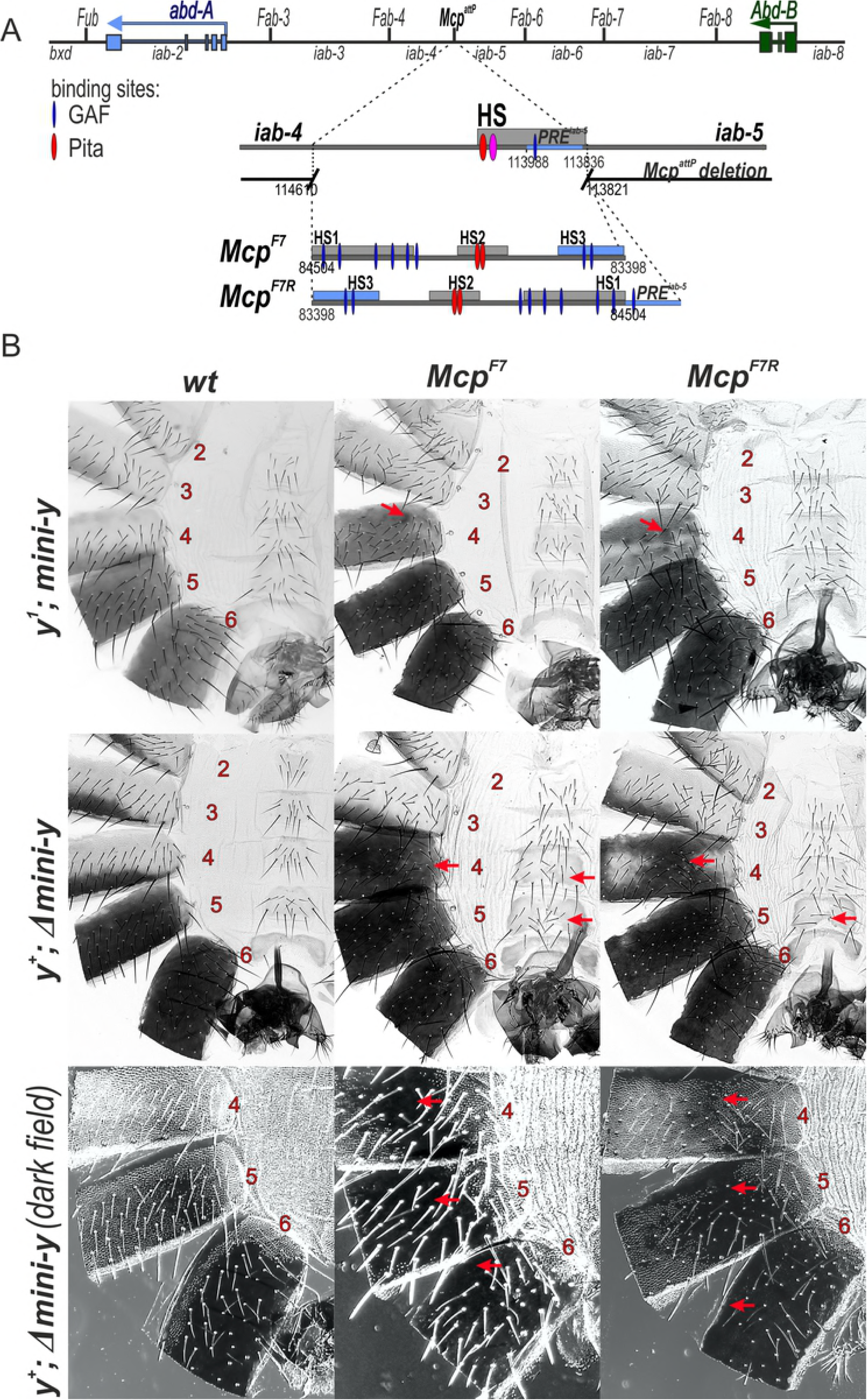
*Mcp*^*F7*^ and *Mcp*^*F7R*^ support *Abd-B* activation in the A4 segment. (A) Schematic representation of the *Fab-7* boundary. The 1.1 kb *Fab-7* replacement consists of HS1, HS2 and HS3 (*iab-7* PRE) regions (shown as gray boxes). (B) Morphology of the 2^th^ to 6^th^ abdominal segments in *Mcp*^*F7*^ and *Mcp*^*F7R*^ males. Other designations are as in Fig.2.

Like the *Mcp*^*CTCF*^ replacement, the *mini-y* reporter is turned on in A5 (PS10) and A6 (PS11) in *Mcp*^*F7*^ males, and the tergites in both of these segments are black. However, *Mcp*^*F7*^ differs in two important respects from *Mcp*^*CTCF*^. First, there are one or two small patches of darkly pigmented cuticle in the A4 tergite (marked by the arrow). These patches are variable and appear to be clonal in origin. This finding indicates that the blocking activity of *Mcp*^*F7*^ is incomplete, and the *mini-y* reporter is ectopically activated by the *iab-4* domain in a small number of PS9 cells. Second, instead of a stripe of light yellow-brown pigmentation along the posterior margin, nearly the entire A4 tergite is covered in yellow-brown pigmentation. This pattern of pigmentation is not observed in A4 in *y*^*1*^ males carrying the *Mcp*^*CTCF*^ replacement (Fig 2) and the *mini-y* reporter or for that matter in wild type *y*^*1*^ males (see Fig 4). The presence of yellow-brown pigmentation throughout most of the A4 tergite suggests that the cells in this segment (PS9) are not properly specified. This is the case. When the *mini-y* reporter was excised and replaced by the endogenous X-linked *y*^*+*^ gene, the A4 tergite has a black pigmentation like A5 and A6 (Fig 4). Since expression of the *yellow* gene is controlled by *Abd-B*, this observation indicates that *Abd-B* must be ectopically activated throughout A4. Antibody staining experiments of the CNS in *Mcp*^*F7*^ embryos indicate that this inference is correct (Fig 3B).

A simple interpretation of these findings is that *Mcp*^*F7*^ is unable to block crosstalk between *iab-4* and *iab-5* and, as a result, *iab-5* is ectopically activated in all PS9 cells. However, such interpretation is inconsistent with the expression pattern of the *mini-y* reporter; it is only activated in small clones in the A4 tergite and not in the entire A4 tergite. By way of comparison, the black pigmentation generated by the reporter in *Mcp*^*PRE*^, which has no boundary activity, was clearly quite different from the light yellow-brown pigmentation observed for the reporter in *Mcp*^*F7*^.

There are other reasons to think that this simple interpretation is incorrect and that *Mcp*^*F7*^ replacement has a much more complicated effect on the operation of *iab-4* and of the regulatory domains that control *Abd-B* expression. In wild type males, the A6 sternite has a banana shape and no bristles, while the A5 and A4 sternites resemble isosceles trapezoids and are covered with bristles. As can be seen in Fig 4, the A4 and A5 sternites in *Mcp*^*F7*^ males are split into two connected lobes and resemble the banana shape of the A6 sternite. These morphological changes are indicative of a GOF transformation of both A4 (PS9) and A5 (PS10) toward an A6 (PS11) identity. This type of transformation is not observed in *Mcp* boundary deletions, nor is it observed in the *Mcp*^*PRE*^ replacement.

Further evidence of a novel A4/A5→A6 transformation can be seen in the pattern of trichome hairs in the tergites. In wild type flies, the A4 and A5 tergites are covered with trichomes, while trichomes are only found along the anterior and ventral margins of the A6 tergite (see darkfield image in Fig 4). In the *Mcp*^*F7*^ replacement, there are large regions of the A4 and A5 tergite that are devoid of trichomes. There are even anomalies in A6: the band of trichomes along the anterior margin is absent. Similar alterations in cuticular phenotypes are observed in *Mcp*^*F7*^ females (S2 Fig). These findings indicate that the normal regulation of *Abd-B* is disrupted in several parasegments when *Mcp* is replaced by the *Fab-7* boundary.

In its endogenous context, the functioning of *Fab-7* is weakly orientation dependent. For this reason, we anticipated that the reverse *Mcp* replacement, *Mcp*^*F7R*^, would alter the *Abd-B* expression pattern in several parasegments and give a similar though perhaps milder spectrum of phenotypic effects. Fig 4 shows that this is the case. In *y*^*+*^ background, large regions of the A4 tergite have a black pigmentation like A5 and A6. The ectopic activation appears to be weaker than in the *Mcp*^*F7*^ replacement as there are regions in A4 in which the endogenous *yellow* gene is not turned on. Also, and unlike *Mcp*^*F7*^, there are no bald patches in the A4 trichomes, while the sternite appears to have a normal isosceles trapezoid shape. However, the novel transformations seen in *Mcp*^*F7*^ in the more posterior segments A5 (PS10) and A6 (PS11) are still evident. The A5 tergite is not completely covered with trichomes, while the trichomes along the anterior margin of A6 are absent. The A5 sternite is also misshapen. Thus, like *Mcp*^*F7*^, introducing a reversed *Fab-7* boundary in place of *Mcp* disrupts *Abd-B* regulation across several parasegments.

### The *Fab-8* boundary displays orientation-dependent effects on ectopic activation of *Abd-B* in the A4 abdominal segment

In previous *Fab-7* replacement experiments we found that a 337 bp fragment (*F8*^*337*^) spanning the *Fab-8* boundary nuclease hypersensitive site is sufficient to fully rescue a *Fab-7* boundary deletion (Kyrchanova et al. 2016). In the direct (forward) orientation this fragment not only blocks crosstalk but also supports bypass. However, when the orientation of the *Fab-8* boundary is reversed, bypass activity is lost, while blocking is unaffected. Since *F8*^*337*^ appears to be fully functioning, we inserted this fragment in both orientations next to the *iab-5 PRE* in the *Mcp* deletion (*Mcp*^*F8*^ and *Mcp*^*F8R*^).

The effects of the *Fab-8* replacement in the reverse orientation, *Mcp*^*F8R*^, will be considered first. Like the *Mcp*^*CTCF*^ replacement, *Mcp*^*F8R*^ blocks crosstalk between *iab-4* and *iab-5* and the *mini-y* reporter is off in A4 (Fig 5). After the deletion of the *mini-y* reporter and introducing a wild type *y*^*+*^ allele, the pigmentation in the adult male abdomen is equivalent to that in wild type flies. The morphological features of *Mcp*^*F8R*^ tegites and sternites also resemble those in wild type flies or the *Mcp*^*CTCF*^ replacement and there is no indication of the other abdominal transformations seen in the *Fab-7* replacements. Consistent with the phenotype of the adult cuticle, the pattern of *Abd-B* expression in the embryonic CNS resembles wild type (Fig 3B). Thus, the *Mcp*^*F8R*^ replacement fully substitutes for the endogenous *Mcp* boundary.

**Fig 5.**
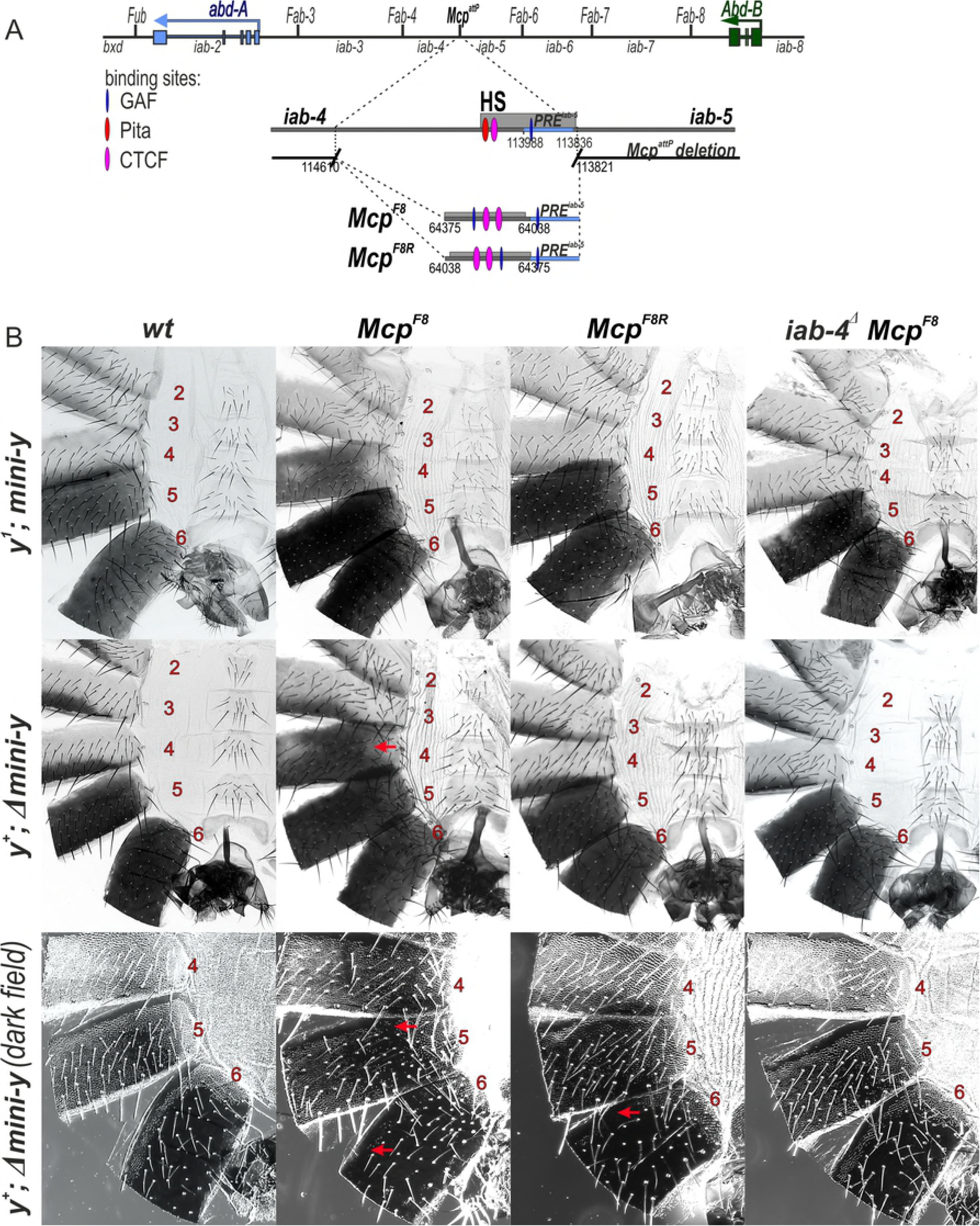
Activation of *Abd-B* by the *iab-4* enhancer depends on the orientation of the *Fab-8* insulator in *Mcp*^*F8*^ and *Mcp*^*F8R*^ mutants. (A) Molecular maps of the *Fab-8* boundary and *F8*^*337*^. The *Fab-8* insulator is shown as a horizontal bar. The proximal and distal deficiency endpoints of the *Fab-8* deletions are shown below. For other designations see Fig 2. (B) Morphology of the 2^nd^ to 6^th^ abdominal segments in insulator in *Mcp*^*F8*^, *Mcp*^*F8R*^ and *iab-4*^*Δ*^ *Mcp*^*F8*^ males. Other designations are as in Fig 2.

A different result is obtained when the *F8*^*337*^ sequence is inserted in its normal forward orientation. Like the reverse orientation *Mcp*^*F8R*^, *Mcp*^*F8*^ efficiently blocks crosstalk between *iab-4* and *iab-5* and the *mini-y* reporter is not activated in A4 (PS9). On the other hand, like the *Fab-7* replacements (*Mcp*^*F7*^ and *Mcp*^*F7R*^) most of the A4 tergite is covered in a light yellow-brown pigmentation instead of the normal stripe of yellow-brown pigmentation along the posterior margin of the tergite that is seen in *y*^*1*^ males. Moreover, when the reporter is excised and the *y*^*1*^ allele replaced by the wild type *y*^*+*^ gene, nearly the entire A4 tergite is black. Consistent with the induction of *y*^*+*^ expression in A4, *Abd-B* is active in PS9 in the embryonic CNS (Fig 3B). The GOF transformation of A4 (PS9)→A5 (PS10) is not the only anomaly in *Mcp*^*F8*^ flies. While there does not seem to be any misspecification of the tergite or sternites in A5 (PS10), the line of trichomes along the anterior margin of the A6 tergite is disrupted or absent altogether indicating that there are abnormalities in the temporal and/or special pattern of *Abd-B* expression in PS11.

### Ectopic expression of *Abd-B* in A4 (PS9) requires a functional *iab-4* domain

In the *Fab-7* replacement experiments, the relative orientation of the *Fab-8* boundary was thought to be important because it determined whether the chromatin loops formed between the replacement boundary and the *AB-I* element and/or the PTE sequence upstream of the *Abd-B* transcription start site were circle loops or stem loops [30,74]. In the forward orientation circle loops are expected to be formed and in this configuration, the downstream *iab-5* regulatory domain is brought into close proximity with the *Abd-B* promoter. In the reverse orientation, *iab-6* and *iab-7* are predicted to form stem loops, and this configuration would tend to isolate the *iab-5* regulatory domain from the *Abd-B* promoter.

It seemed possible that a similar mechanism might be in play in the *Fab-8* replacements of *Mcp*. In the forward orientation (*Mcp*^*F8*^), the *iab-4* regulatory domain would be brought into close proximity to the *Abd-B* gene, activating its ectopic expression in A4 (PS9). In the opposite orientation, the spatial relationship between the *iab-4* domain and the *Abd-B* promoter would not be conducive for activation. In this case, *Abd-B* would be off in A4 (PS9). A strong prediction of this model is that the inappropriate activation of *Abd-B* in PS9 in the *Mcp*^*F8*^ replacement should depend on a functional *iab-4* domain.

To test this prediction, we used CRISPR (see S3 Fig) to delete a 4,401 bp sequence (*iab-4*^*Δ*^) that spans the putative *iab-4* initiation element in flies carrying the *Mcp*^*F8*^ replacement. The *iab-4*^*Δ*^ sequence was selected based on the clustering of multiple binding sites for embryonic gap and pair-rule gene proteins [75]. Fig 5 shows that the ectopic activation of *y*^*+*^ in A4 in *Mcp*^*F8*^ flies was eliminated by the *iab-4*^*Δ*^ deletion. Moreover, and as predicted, *Abd-B* was not activated in A4 (PS9) in the embryonic CNS of *iab-4*^*Δ*^ *Mcp*^*F8*^ embryos (Fig 3B). Interestingly, the loss of trichomes along the anterior margin of the A6 tergite in *Mcp*^*F8*^ also seemed to depend on a functional *iab-4* domain. As can be seen in Fig 5, the trichome pattern in the A6 tergite of *iab-4*^*Δ*^ *Mcp*^*F8*^ flies resembled that of wild type.

## Conclusion

*Mcp* defines the boundary between the regulatory domains that control expression of *abd-A* and *Abd-B*. In this location, it is required to block crosstalk between the flanking domains *iab-4* and *iab-5*, but it does not need to mediate bypass. In this respect, it differs from the boundaries that are located within the set of regulatory domains that control either *abd-A* or *Abd-B*, as these boundaries must have both activities. Consistent with this limited role, we found that *Mcp* can be replaced by multimerized binding sites for the dCTCF protein. Quite different results are obtained when *Mcp* is replaced by *Fab-7* or *Fab-8*. Although *Fab-7* is able to block crosstalk between *iab-4* and *iab-5*, its blocking activity is incomplete and there are small clones of cells in which the *mini-y* reporter is activated in A4. In contrast, the *mini-y* reporter is off throughout A4 in the *Fab-8* boundary replacements, indicating that it efficiently blocks crosstalk between *iab-4* and *iab-5*. One plausible reason for this difference is that *Mcp* and the boundaries flanking *Mcp* (*Fab-4* and *Fab-6*) utilize dCTCF as does *Fab-8*, while this architectural protein does not bind to *Fab-7* [33].

In spite of their normal (or near normal) blocking activity, both boundaries still perturb *Abd-B* regulation. In the case of *Fab-8*, the misregulation of *Abd-B* is orientation dependent just like the bypass activity of this boundary when it is used to replace *Fab-7* [30]. When inserted in the reverse orientation, *Fab-8* behaves like multimerized dCTCF sites and it fully rescues the *Mcp* deletion. In contrast, when inserted in the forward orientation, *Fab-8* induces the expression of *Abd-B* in A4 (PS9), and the misspecification of this parasegment. Our results, taken together with our previous studies [30,59,60], support a model in which the chromatin loops formed by *Fab-8* inserted at *Mcp* in the forward orientation brings the enhancers in the *iab-4* regulatory domain in close proximity to the *Abd-B* promoter, leading to the activation of *Abd-B* in A4 (PS9). In contrast, when inserted in the opposite orientation, the chromatin loops formed by the ectopic *Fab-8* boundary are not permissive for interactions between *iab-4* and the *Abd-B* promoter. Importantly, the ectopic activation of *Abd-B* in A4 when *Fab-8* is inserted in the forward orientation suggests that the bypass activity has a predisposed preference, namely it is targeted for interactions with the *Abd-B* gene. From this perspective, it would appear that boundary bypass for the regulatory domains that control *Abd-B* expression is not a passive process in which the boundaries are simply permissive for interactions between the regulatory domains and the *Abd-B* promoter. Instead, it appears to be an active process in which the boundaries are responsible for bringing the regulatory domains into contact with the *Abd-B* gene. It clearly will be of interest to test the out of context functional properties of the boundaries associated with the *abd-B* and *Ubx* genes to see if they behave like *Fab-7* and *Fab-8*.

While similar conclusions can be drawn from the induction of *Abd-B* expression in A4 (PS9) when *Fab-7* is inserted in place of *Mcp*, this boundary causes even more profound disruptions in the normal pattern of *Abd-B* regulation. In the forward orientation, A4 and A5 are transformed towards an A6 identity, while A6 is also misspecified. Similar though somewhat less severe effects are observed when *Fab-7* is inserted in the reverse orientation. Although the mechanisms responsible for these novel phenotypic effects are uncertain, a plausible idea is that pairing interactions between the *Fab-7* insert and the endogenous *Fab-7* boundary disrupt the normal topological organization of the regulatory domains in a manner similar to that seen in boundary competition transgene assays [76]. Further studies will be required to test this idea.

## Materials and Methods

### Generation of *Mcp*^*attP*^ by *CRISPR/Cas9*-induced homologous recombination

The backbone of the recombination plasmid was designed *in silico* and contains several genetic elements in the following order: [MCS5]-[attP]-[3xP3-EGFP-SV40polyA]-[attP]-[FRT]-[MCS3]. This DNA fragment was synthesized and cloned into pUC57 by Genewiz. The two multiple cloning sites MCS5 and MCS3 were used to clone homology arms into this plasmid. The orientations of two the *attP* sites are inverted relative to each other and serve as targets for *ɸC31*-mediated recombination mediated cassette exchange [77]. The *3×3P-EGFP* reporter [78] was introduced as a means to isolate positive recombination events. The *Flp*-recombinase target *FRT* [79] were includedl for the deletion of the selectable *mini-yellow* marker after recombination mediated cassette exchange.

Homology arms were PCR-amplified from *y w* genomic DNA using the following primers: CCTGCCGACTGAACGAATGC and ACGCCCTGATCCCGATACACATAC for the proximal arm (*iab-4* side; 3967 bp fragment), and GCGTTTGTGTGTAGTAAATGTATC and AAAGGCCAACAAAGAACACATGGACG for the distal arm (*iab-5* side; 4323 bp fragment). A successful homologous recombination event will generate a 789 bp deletion within the *Mcp* region (Genome Release R6.22: 3R:16’868’830 – 16’869’619; or complete sequence of BX-C (Martin et al. 1995): 113821 - 114610).

The recombination plasmid was injected into *y w vas-Cas9* embryos together with two gRNAs containing the following guides: GCTGGCTTTTACAGCATTTC and GCTTTGTTACCCCTGAAAAT. Concentrations were as described in Gratz et al.[80]. The injected embryos were grown to adulthood and crossed with *y w* partners. Among the few fertile crosses, one produced many larvae with a clear GFP-signal in the posterior part of their abdomens. This observation suggested that these animals had successfully integrated the recombination plasmid and that the *3×3P-EGFP* reporter acts as an enhancer trap for *Abd-B* specific enhancers. GFP positive larvae were isolated and grown to adulthood. Emerging males showed the expected *Mcp* phenotype. Also, and as expected for a reporter located in the BX-C, no fluorescence signal could be detected in their eyes, indicating that the *3×3P-EGFP* reporter is silenced in eye cells where the *3×3P* promoter is usually active. The planned homologous recombination event could later be verified by PCR and sequencing. We will refer to it as *Mcp*^*attP*^.

12 *EGFP*-and *Mcp*-positive candidate males were individually crossed with *y w* virgins. Only 2 were fertile. The sterility of others may be caused by presence of *off*-targets as afrequent non-specific result of CRISPR/*Cas9* editing. Starting from the two fertile crosses, 2 independent balanced stocks could be obtained according to established crossing schemes. One of them was used to obtain a *y w M{vas-integrase}zh-2A; Mcp*^*attP*^*/TM3,Sb* stock for recombination mediated cassette exchange. Because of poor survival rates in injection experiments, the *Mcp*^*attP*^ chromosome was also temporarily combined with *Dp(3R)P9, Sb* (*y w M{vas-integrase}zh-2A; Mcp*^*attP*^*/Dp(3R)P9, Sb*). By selection we obtained homozygous *Mcp*^*attP*^ line that was subsequently used for fly injections.

### Generation of *iab-4*^*Δ*^ by CRISPR/Cas9-induced homologous recombination

For generating dsDNA donors for homology-directed repair we used *pHD-DsRed* vector that was a gift from Kate O’Connor-Giles (Addgene plasmid # 51434). The final plasmid contains genetic elements in the following order: [*iab-4* proximal arm]-[attP]-[lox]-[3xP3-dsRed-SV40polyA]-[lox]-[*iab-4* distal arm]. Homology arms were PCR-amplified from *yw* genomic DNA using the following primers: TTT*GAATTC*TTCCAGACACGCATCGGG and AAA*CATATG*CTTGCTATCGACCCTCCTC for the proximal arm (846 bp fragment), and AAT*ACTAGT*CTCGGAAAGGGAAGAAGTTC and TAC*TCGAGC*CGCTAAAGGACGTTCTGC for the distal arm (835 bp fragment). A successful homologous recombination event will generate a 4401 bp deletion within the *iab-4* region (Genome Release R6.22: 3R:16,861,368..16,869,768; or complete sequence of BX-C [4]: 120073-115673).

Targets for *Cas9* were selected using “CRISPR optimal target finder” – the program from O’Connor-Giles Lab. The recombination plasmid was injected into *Mcp*^*F8*^ *vasa-Cas9* embryos together with two gRNAs containing the following guides: ATAGCAAGTAGGAGTGGAGT and GAACTTCTTCCCTTTCCGAGCGG. Concentrations were as described in Gratz et al. (2014). Injectees were grown to adulthood and crossed with *y w*; *TM6/MKRS* partners. Flies with clear dsRed-signal in eyes and the posterior part of their abdomens were selected into a new separate line. The successfully integration of the recombination plasmid was verified by PCR.

### Cuticle preparations

Adult abdominal cuticles of homozygous enclosed 3-4 day old flies were prepared essentially as described in (Kyrchanova et al. 2017) and mounted in 100% glycerol. Photographs in the bright or dark field were taken on the Nikon SMZ18 stereomicroscope using Nikon DS-Ri2 digital camera, processed with ImageJ 1.50c4 and Fiji bundle 2.0.0-rc-46.

### Embryo immunostaining

Primary antibodies were mouse monoclonal anti-Abd-B at 1:100 dilution (1A2E9, generated by S.Celniker, deposited to the Developmental Studies Hybridoma Bank) and polyclonal rabbit anti-Engrailed at 1:1000 dilution (a kind gift from Judith Kassis). Secondary antibodies were goat anti-mouse Alexa Fluor 488 and anti-rabbit Alexa Fluor 647 (Molecular Probes) at 1:2000 dilution. Stained embryos were mounted in the following solution: 23% glycerol, 10% Mowiol 4-88, 0.1M Tris-HCl pH 8.3. Images were acquired on Leica TCS SP-2 confocal microscope and processed using GIMP 2.8.16, ImageJ 1.50c4, Fiji bundle 2.0.0-rc-46.

## Acknowledgments

We thank Farhod Hasanov and Aleksander Parshikov for fly injections. This study was performed using the equipment of the IGB RAS facilities supported by the Ministry of Science and Education of the Russian Federation.

## Supporting information captions

**S1 Fig. The strategy to create *Mcp* replacement lines.** On the top: schematic representation of regulatory region of the *abd-A* and *Abd-B* genes (blue and green, respectively). The 789 bp *Mcp* region that was deleted (coordinates according to complete sequence of BX-C in SEQ89E numbering) and replaced by two *attP* sites for the integration of the tested constructs. *3xP3-eGFP* was used as a marker gene. *frt* site was used for excision of *yellow* maker gene. The plasmid that was injected into *Mcp*^*attP*^ line, contains two *attB* site for integration, *iab-5* PRE for restoring functional integrity of the *iab-5* domain, the *frt* site for excision of *yellow* gene, *lox* sites for excision of testing element. Testing elements were inserted just in front of *iab-5* PRE.

**S2 Fig. The abdominal cuticles of *wt, Mcp*^*F8*^, *Mcp*^*F8R*^, *Mcp*^*F7*^ and *Mcp*^*F7R*^ females.** Morphology of the 2^th^ to 6^th^ abdominal segments in *wt, Mcp*^*F8*^, *Mcp*^*F8R*^, *Mcp*^*F7*^ and *Mcp*^*F7R*^ females. The expression of *mini-y* (black pigment) is shown on the upper panel. Localization of trichomes on tergites is shown lower.

**S3 Fig. The strategy to create *iab-4* deletion.** The scheme of the regulatory region in the distal part of the BX-C. Horizontal arrows represent transcripts for *abd-A* (blue) and *Abd-B* (green). The *iab-4* region was selected using FlyBase, based on the clustering of multiple binding sites for embryonic gap and pair-rule gene proteins. The screenshot show localization of the 4401 bp of *iab-4* deletion with R6 genome release coordinates. The coordinates of *iab-4* deletion according to complete sequence of BX-C (in SEQ89E numbering) are 120073-115673 (shown lower). The deletion was made using CRISPR/Cas9 strategy. Targets for Cas9 were selected using “CRISPR optimal target finder” – program from O’Connor-Giles Lab. Vector for generating dsDNA donors for homology-directed repair contains the visible marker 3xP3-DsRed. pHD-DsRed was a gift from Kate O’Connor-Giles (Addgene plasmid # 51434). *dsRed* gene was using for selection of flies with *iab-4* deletion.

## References

1. Lewis EB. A gene complex controlling segmentation in Drosophila. Nature. 1978/12/07. 1978;276: 565–570. Available: http://www.ncbi.nlm.nih.gov/pubmed/103000

2. Mihaly J, Hogga I, Barges S, Galloni M, Mishra RK, Hagstrom K, et al. Chromatin domain boundaries in the Bithorax complex. Cell Mol Life Sci. 1998/03/06. 1998;54: 60–70. Available: http://www.ncbi.nlm.nih.gov/pubmed/9487387

3. Sanchez-Herrero E, Vernos I, Marco R, Morata G. Genetic organization of Drosophila bithorax complex. Nature. 1985/01/10. 1985;313: 108–113. Available: http://www.ncbi.nlm.nih.gov/pubmed/3917555

4. Martin CH, Mayeda CA, Davis CA, Ericsson CL, Knafels JD, Mathog DR, et al. Complete sequence of the bithorax complex of Drosophila. Proc Natl Acad Sci U S A. 1995;92: 8398–8402. Available: http://www.ncbi.nlm.nih.gov/pubmed/7667301

5. McCall K, O’Connor MB, Bender W. Enhancer traps in the Drosophila bithorax complex mark parasegmental domains. Genetics. 1994;138: 387–399. Available: http://www.ncbi.nlm.nih.gov/pubmed/7828822

6. Celniker SE, Sharma S, Keelan DJ, Lewis EB. The molecular genetics of the bithorax complex of Drosophila: cis-regulation in the Abdominal-B domain. EMBO J. 1990;9: 4277–4286. Available: http://www.ncbi.nlm.nih.gov/pubmed/2265608

7. Bender W, Akam M, Karch F, Beachy PA, Peifer M, Spierer P, et al. Molecular Genetics of the Bithorax Complex in Drosophila melanogaster. Science (80-). 1983;221: 23–29. doi:10.1126/science.221.4605.23

8. Duncan I. The bithorax complex. Annu Rev Genet. 1987/01/01. 1987;21: 285–319. doi:10.1146/annurev.ge.21.120187.001441

9. Karch F, Bender W, Weiffenbach B. abdA expression in Drosophila embryos. Genes Dev. 1990;4: 1573–1587. doi:10.1101/gad.4.9.1573

10. François Karch et. al. The abdominal region of the bithorax complex. Cell. 1985;43: 81–96. Available: <https://www.ncbi.nlm.nih.gov/pubmed/3935319

11. Boulet AM, Lloyd A, Sakonju S. Molecular definition of the morphogenetic and regulatory functions and the cis-regulatory elements of the Drosophila Abd-B homeotic gene. Development. 1991;111: 393–405.

12. Sánchez-Herrero E. Control of the expression of the bithorax complex genes abdominal-A and abdominal-B by cis-regulatory regions in Drosophila embryos. Development. 1991;111: 437–449.

13. Maeda RK, Karch F. The ABC of the BX-C: the bithorax complex explained. Development. 2006/03/25. 2006;133: 1413–1422. doi:10.1242/dev.02323

14. Maeda RK, Karch F. The open for business model of the bithorax complex in Drosophila. Chromosoma. 2015;124: 293–307. doi:10.1007/s00412-015-0522-0

15. Maeda RK, Karch F. Cis-regulation in the Drosophila Bithorax Complex. Adv Exp Med Biol. 2010/08/28. 2010;689: 17–40. Available: http://www.ncbi.nlm.nih.gov/pubmed/20795320

16. Maeda RK, Karch F. Gene expression in time and space: additive vs hierarchical organization of cis-regulatory regions. Curr Opin Genet Dev. 2011/02/26. 2011;21: 187–193. doi:10.1016/j.gde.2011.01.021

17. Kassis JA, Brown JL. Polycomb group response elements in Drosophila and vertebrates. Adv Genet. 2013/02/20. 2013;81: 83–118. doi:10.1016/B978-0-12-407677-8.00003-8

18. Mihaly J, Hogga I, Gausz J, Gyurkovics H, Karch F. In situ dissection of the Fab-7 region of the bithorax complex into a chromatin domain boundary and a Polycomb-response element. Development. 1997/05/01. 1997;124: 1809–1820. Available: http://www.ncbi.nlm.nih.gov/pubmed/9165128

19. Mihaly J, Barges S, Sipos L, Maeda R, Cleard F, Hogga I, et al. Dissecting the regulatory landscape of the Abd-B gene of the bithorax complex. Development. 2006/07/05. 2006;133: 2983–2993. doi:10.1242/dev.02451

20. Barges S, Mihaly J, Galloni M, Hagstrom K, Muller M, Shanower G, et al. The Fab-8 boundary defines the distal limit of the bithorax complex iab-7 domain and insulates iab-7 from initiation elements and a PRE in the adjacent iab-8 domain. Development. 2000/01/29. 2000;127: 779–790. Available: http://www.ncbi.nlm.nih.gov/pubmed/10648236

21. Gyurkovics H, Gausz J, Kummer J, Karch F. A new homeotic mutation in the Drosophila bithorax complex removes a boundary separating two domains of regulation. EMBO J. 1990/08/01. 1990;9: 2579–2585. Available: http://www.ncbi.nlm.nih.gov/pubmed/1973385

22. Galloni M, Gyurkovics H, Schedl P, Karch F. The bluetail transposon: evidence for independent cis-regulatory domains and domain boundaries in the bithorax complex. EMBO J. 1993;12: 1087–1097. Available: http://www.ncbi.nlm.nih.gov/pubmed/8384551

23. Karch F, Galloni M, Sipos L, Gausz J, Gyurkovics H, Schedl P. Mcp and Fab-7: molecular analysis of putative boundaries of cis-regulatory domains in the bithorax complex of Drosophila melanogaster. Nucleic Acids Res. 1994/08/11. 1994;22: 3138–3146. Available: http://www.ncbi.nlm.nih.gov/pubmed/7915032

24. Bender W, Lucas M. The border between the ultrabithorax and abdominal-A regulatory domains in the Drosophila bithorax complex. Genetics. 2013;193: 1135–1147. doi:10.1534/genetics.112.146340

25. Bowman SK, Deaton AM, Domingues H, Wang PI, Sadreyev RI, Kingston RE, et al. H3K27 modifications define segmental regulatory domains in the Drosophila bithorax complex. Elife. 2014;3: e02833. doi:10.7554/eLife.02833

26. Iampietro C, Cleard F, Gyurkovics H, Maeda RK, Karch F. Boundary swapping in the Drosophila Bithorax complex. Development. 2008/11/07. 2008;135: 3983–3987. doi:10.1242/dev.025700

27. Iampietro C, Gummalla M, Mutero A, Karch F, Maeda RK. Initiator elements function to determine the activity state of BX-C enhancers. PLoS Genet. 2011/01/05. 2010;6: e1001260. doi:10.1371/journal.pgen.1001260

28. Kyrchanova O, Mogila V, Wolle D, Magbanua JP, White R, Georgiev P, et al. The boundary paradox in the Bithorax complex. Mech Dev. 2015;138: 122–132. doi:10.1016/j.mod.2015.07.002

29. Hogga I, Mihaly J, Barges S, Karch F. Replacement of Fab-7 by the gypsy or scs insulator disrupts long-distance regulatory interactions in the Abd-B gene of the bithorax complex. Mol Cell. 2001;8: 1145–1151. Available: http://www.ncbi.nlm.nih.gov/pubmed/11741549

30. Kyrchanova O, Mogila V, Wolle D, Deshpande G, Parshikov A, Cléard F, et al. Functional Dissection of the Blocking and Bypass Activities of the Fab-8 Boundary in the Drosophila Bithorax Complex. PLoS Genet. 2016;12: e1006188. doi:10.1371/journal.pgen.1006188

31. Kyrchanova O, Zolotarev N, Mogila V, Maksimenko O, Schedl P, Georgiev P. Architectural protein Pita cooperates with dCTCF in organization of functional boundaries in Bithorax complex. Development. 2017;144: 2663–2672. doi:10.1242/dev.149815

32. Maksimenko O, Bartkuhn M, Stakhov V, Herold M, Zolotarev N, Jox T, et al. Two new insulator proteins, Pita and ZIPIC, target CP190 to chromatin. Genome Res. 2014/10/25. 2015;25: 89–99. doi:10.1101/gr.174169.114

33. Holohan EE, Kwong C, Adryan B, Bartkuhn M, Herold M, Renkawitz R, et al. CTCF genomic binding sites in Drosophila and the organisation of the bithorax complex. PLoS Genet. 2007/07/10. 2007;3: e112. doi:10.1371/journal.pgen.0030112

34. Magbanua JP, Runneburger E, Russell S, White R. A Variably Occupied CTCF Binding Site in the Ultrabithorax Gene in the Drosophila Bithorax Complex. Mol Cell Biol. 2015;35: 318–330. doi:10.1128/MCB.01061-14

35. Moon H, Filippova G, Loukinov D, Pugacheva E, Chen Q, Smith ST, et al. CTCF is conserved from Drosophila to humans and confers enhancer blocking of the Fab-8 insulator. EMBO Rep. 2005/01/29. 2005;6: 165–170. doi:10.1038/sj.embor.7400334

36. Gruzdeva N, Kyrchanova O, Parshikov A, Kullyev A, Georgiev P. The Mcp element from the bithorax complex contains an insulator that is capable of pairwise interactions and can facilitate enhancer-promoter communication. Mol Cell Biol. 2005/04/16. 2005;25: 3682–3689. doi:10.1128/MCB.25.9.3682-3689.2005

37. Schweinsberg S, Hagstrom K, Gohl D, Schedl P, Kumar RP, Mishra R, et al. The enhancer-blocking activity of the Fab-7 boundary from the Drosophila bithorax complex requires GAGA-factor-binding sites. Genetics. 2004/12/08. 2004;168: 1371–1384. doi:10.1534/genetics.104.029561

38. Schweinsberg SE, Schedl P. Developmental modulation of Fab-7 boundary function. Development. 2004/08/27. 2004;131: 4743–4749. doi:10.1242/dev.01343

39. Perez-Lluch S, Cuartero S, Azorin F, Espinas ML. Characterization of new regulatory elements within the Drosophila bithorax complex. Nucleic Acids Res. 2008;36: 6926–6933. doi:10.1093/nar/gkn818

40. Zhou J, Ashe H, Burks C, Levine M. Characterization of the transvection mediating region of the abdominal-B locus in Drosophila. Development. 1999/06/22. 1999;126: 3057–3065. Available: http://www.ncbi.nlm.nih.gov/pubmed/10375498

41. Zhou J, Barolo S, Szymanski P, Levine M. The Fab-7 element of the bithorax complex attenuates enhancer-promoter interactions in the Drosophila embryo. Genes Dev. 1996/12/15. 1996;10: 3195–3201. Available: http://www.ncbi.nlm.nih.gov/pubmed/8985187

42. Hagstrom K, Muller M, Schedl P. Fab-7 functions as a chromatin domain boundary to ensure proper segment specification by the Drosophila bithorax complex. Genes Dev. 1996/12/15. 1996;10: 3202–3215. Available: http://www.ncbi.nlm.nih.gov/pubmed/8985188

43. Zhou J, Levine M. A novel cis-regulatory element, the PTS, mediates an anti-insulator activity in the Drosophila embryo. Cell. 1999/12/28. 1999;99: 567–575. Available: http://www.ncbi.nlm.nih.gov/pubmed/10612393

44. Chen Q, Lin L, Smith S, Lin Q, Zhou J. Multiple Promoter Targeting Sequences exist in Abdominal-B to regulate long-range gene activation. Dev Biol. 2005;286: 629–636. doi:10.1016/j.ydbio.2005.08.025

45. Lin Q.; Wu D.; Zhou J. The promoter targeting sequence facilitates and restricts a distant enhancer to a single promoter in the Drosophila embryo. Development. 2003;130: 519–526.

46. Cai HN, Shen P. Effects of cis arrangement of chromatin insulators on enhancer-blocking activity. Science (80- ). 2001/02/13. 2001;291: 493–495. doi:10.1126/science.291.5503.493

47. Muravyova E, Golovnin A, Gracheva E, Parshikov A, Belenkaya T, Pirrotta V, et al. Loss of insulator activity by paired Su(Hw) chromatin insulators. Science (80- ). 2001/02/13. 2001;291: 495–498. doi:10.1126/science.291.5503.495

48. Sigrist CJ, Pirrotta V. Chromatin insulator elements block the silencing of a target gene by the Drosophila polycomb response element (PRE) but allow trans interactions between PREs on different chromosomes. Genetics. 1997/09/01. 1997;147: 209–221. Available: http://www.ncbi.nlm.nih.gov/pubmed/9286681

49. Muller M, Hagstrom K, Gyurkovics H, Pirrotta V, Schedl P. The mcp element from the Drosophila melanogaster bithorax complex mediates long-distance regulatory interactions. Genetics. 1999/11/05. 1999;153: 1333–1356. Available: http://www.ncbi.nlm.nih.gov/pubmed/10545463

50. Li HB, Muller M, Bahechar IA, Kyrchanova O, Ohno K, Georgiev P, et al. Insulators, not Polycomb response elements, are required for long-range interactions between Polycomb targets in Drosophila melanogaster. Mol Cell Biol. 2010/12/08. 2011;31: 616–625. doi:10.1128/MCB.00849-10

51. Fujioka M, Mistry H, Schedl P, Jaynes JB. Determinants of Chromosome Architecture: Insulator Pairing in cis and in trans. PLoS Genet. 2016;2: e1005889. doi:10.1371/journal.pgen.1005889

52. Kyrchanova O, Chetverina D, Maksimenko O, Kullyev A, Georgiev P. Orientation-dependent interaction between Drosophila insulators is a property of this class of regulatory elements. Nucleic Acids Res. 2008/11/07. 2008;36: 7019–7028. doi:10.1093/nar/gkn781

53. Hendrickson JE, Sakonju S. Cis and trans interactions between the iab regulatory regions and abdominal-A and abdominal-B in Drosophila melanogaster. Genetics. 1995;139: 835–848.

54. Hopmann R, Duncan D, Duncan I. Transvection in the iab-5,6,7 region of the bithorax complex of Drosophila: Homology independent interactions in trans. Genetics. 1995;139: 815–833.

55. Sipos L, Mihály J, Karch F, Schedl P, Gausz J, Gyurkovics H. Transvection in the Drosophila Abd-B domain: Extensive upstream sequences are involved in anchoring distant cis-regulatory regions to the promoter. Genetics. 1998;149: 1031–1050.

56. Ho MC, Schiller BJ, Akbari OS, Bae E, Drewell RA. Disruption of the Abdominal-B promoter tethering element results in a loss of long-range enhancer-directed Hox gene expression in Drosophila. PLoS One. 2011;6: e16283. doi:10.1371/journal.pone.0016283

57. Akbari OS, Schiller BJ, Goetz SE, Ho MCW, Bae E, Drewell RA. The Abdominal-B promoter tethering element mediates promoter-enhancer specificity at the Drosophila bithorax complex. Fly (Austin). 2007;1: 337–339. doi:10.4161/fly.5607

58. Akbari OS, Bae E, Johnsen H, Villaluz A, Wong D, Drewell RA. A novel promoter-tethering element regulates enhancer-driven gene expression at the bithorax complex in the Drosophila embryo. Development. 2007;135: 123–131. doi:10.1242/dev.010744

59. Kyrchanova O, Toshchakov S, Podstreshnaya Y, Parshikov A, Georgiev P. Functional Interaction between the Fab-7 and Fab-8 Boundaries and the Upstream Promoter Region in the Drosophila Abd-B Gene. Mol Cell Biol. 2008;28: 4188–4195. doi:10.1128/MCB.00229-08

60. Kyrchanova O, Ivlieva T, Toshchakov S, Parshikov A, Maksimenko O, Georgiev P. Selective interactions of boundaries with upstream region of Abd-B promoter in Drosophila bithorax complex and role of dCTCF in this process. Nucleic Acids Res. 2010/12/15. 2011;39: 3042–3052. doi:10.1093/nar/gkq1248

61. Busturia a, Lloyd A, Bejarano F, Zavortink M, Xin H, Sakonju S. The MCP silencer of the Drosophila Abd-B gene requires both Pleiohomeotic and GAGA factor for the maintenance of repression. Development. 2001;128: 2163–2173.

62. Bischof J, Maeda RK, Hediger M, Karch F, Basler K. An optimized transgenesis system for Drosophila using germ-line-specific phiC31 integrases. Proc Natl Acad Sci U S A. 2007/03/16. 2007;104: 3312–3317. doi:10.1073/pnas.0611511104

63. Gibert JM, Peronnet F, Schlötterer C. Phenotypic plasticity in Drosophila pigmentation caused by temperature sensitivity of a chromatin regulator network. PLoS Genet. 2007;3: e30. doi:10.1371/journal.pgen.0030030

64. Jeong S, Rokas A, Carroll SB. Regulation of Body Pigmentation by the Abdominal-B Hox Protein and Its Gain and Loss in Drosophila Evolution. Cell. 2006;125: 1387–1399. doi:10.1016/j.cell.2006.04.043

65. Camino EM, Butts JC, Ordway A, Vellky JE, Rebeiz M, Williams TM. The Evolutionary Origination and Diversification of a Dimorphic Gene Regulatory Network through Parallel Innovations in cis and trans. PLoS Genet. 2015;11: e1005136. doi:10.1371/journal.pgen.1005136

66. Rebeiz M, Williams TM. Using Drosophila pigmentation traits to study the mechanisms of cis-regulatory evolution. Curr Opin Insect Sci. 2017;19: 1–7. doi:10.1016/j.cois.2016.10.002

67. Bender W, Hudson A. P element homing to the Drosophila bithorax complex. Development. 2000;127: 3981–3992.

68. Bonchuk A, Maksimenko O, Kyrchanova O, Ivlieva T, Mogila V, Deshpande G, et al. Functional role of dimerization and CP190 interacting domains of CTCF protein in Drosophila melanogaster. BMC Biol. 2015; doi:10.1186/s12915-015-0168-7

69. Fedotova AA, Bonchuk AN, Mogila VA, Georgiev PG. C2H2 zinc finger proteins: The largest but poorly explored family of higher eukaryotic transcription factors. Acta Naturae. 2017;9: 47–58.

70. Wolle D, Cleard F, Aoki T, Deshpande G, Schedl P, Karch F. Functional Requirements for Fab-7 Boundary Activity in the Bithorax Complex. Mol Cell Biol. 2015;35: 3739–3752. doi:10.1128/MCB.00456-15

71. Cleard F, Wolle D, Taverner AM, Aoki T, Deshpande G, Andolfatto P, et al. Different evolutionary strategies to conserve chromatin boundary function in the bithorax complex. Genetics. 2017;205: 589–603. doi:10.1534/genetics.116.195586

72. Kyrchanova, O., Kurbidaeva A, Sabirov M, Postika N, Wolle D, Aoki T, Maksimenko O, Mogila V SP and GP. The Bithorax comlex iab-7 Polycomb Response Element has a novel role in the functioning of the Fab-7 chromatin boundary. Plos Genet. 2018;in press.

73. Mishra RK, Mihaly J, Barges S, Spierer A, Karch F, Hagstrom K, et al. The iab-7 polycomb response element maps to a nucleosome-free region of chromatin and requires both GAGA and pleiohomeotic for silencing activity. Mol Cell Biol. 2001/02/07. 2001;21: 1311–1318. doi:10.1128/MCB.21.4.1311-1318.2001

74. Chetverina D, Fujioka M, Erokhin M, Georgiev P, Jaynes JB, Schedl P. Boundaries of loop domains (insulators): Determinants of chromosome form and function in multicellular eukaryotes. BioEssays. 2017;39. doi:10.1002/bies.201600233

75. Starr MO, Ho MCW, Gunther EJM, Tu YK, Shur AS, Goetz SE, et al. Molecular dissection of cis-regulatory modules at the Drosophila bithorax complex reveals critical transcription factor signature motifs. Dev Biol. 2011;359: 290–302. doi:10.1016/j.ydbio.2011.07.028

76. Gohl D, Aoki T, Blanton J, Shanower G, Kappes G, Schedl P. Mechanism of chromosomal boundary action: roadblock, sink, or loop? Genetics. 2011/01/05. 2011;187: 731–748. doi:10.1534/genetics.110.123752

77. Bateman JR, Lee AM, Wu CT. Site-specific transformation of Drosophila via???C31 integrase-mediated cassette exchange. Genetics. 2006;173: 769–777. doi:10.1534/genetics.106.056945

78. Horn C, Jaunich B, Wimmer EA. Highly sensitive, fluorescent transformation marker for Drosophila transgenesis. Dev Genes Evol. 2000;210: 623–629. doi:10.1007/s004270000111

79. Golic KG, Lindquist S. The FLP recombinase of yeast catalyzes site-specific recombination in the Drosophila genome. Cell. 1989/11/03. 1989;59: 499–509. Available: http://www.ncbi.nlm.nih.gov/pubmed/2509077

80. Gratz SJ, Ukken FP, Rubinstein CD, Thiede G, Donohue LK, Cummings AM, et al. Highly specific and efficient CRISPR/Cas9-catalyzed homology-directed repair in Drosophila. Genetics. 2014;196: 961–971. doi:10.1534/genetics.113.160713

